# *fruitless* mutant male mosquitoes gain attraction to human odor

**DOI:** 10.1101/2020.09.04.282434

**Authors:** Nipun S. Basrur, Maria Elena De Obaldia, Takeshi Morita, Margaret Herre, Ricarda K. von Heynitz, Yael N. Tsitohay, Leslie B. Vosshall

## Abstract

While sexual dimorphism in courtship and copulation behavior is common in the animal kingdom, sexual dimorphism in feeding behavior is rare. The *Aedes aegypti* mosquito provides an example of extreme sexual dimorphism in feeding, because only the females show strong attraction to humans, and bite them to obtain a blood-meal necessary to stimulate egg production^1-8^. The genetic basis of this complex, modular, and sexually dimorphic feeding behavior is unknown. The *fruitless* gene is sex-specifically spliced in the brain of multiple insect species including mosquitoes^9-11^ and encodes a BTB zinc-finger transcription factor that has been proposed to be a master regulator of male courtship and mating behavior across insects^12-17^. Here we use CRISPR-Cas9 to mutate the *fruitless* gene in male mosquitoes. *fruitless* mutant males fail to mate, confirming the ancestral function of this gene in male sexual behavior. Remarkably, *fruitless* mutant males also gain strong attraction to a live human host, a behavior that wild-type males never display. Humans produce multiple sensory cues that attract mosquitoes and we show that *fruitless* specifically controls host-seeking in response to human odor. These results suggest that male mosquitoes possess the neural circuits required to host-seek and that removing *fruitless* reveals this latent behavior in males. Our results highlight an unexpected repurposing of a master regulator of male-specific sexual behavior to control one module of female-specific blood-feeding behavior in a deadly vector of infectious diseases.

## INTRODUCTION

Across animals, males and females of the same species show striking differences in behavior. Male *Paradisaeidae* birds-of-paradise perform an elaborate courtship dance to seduce prospective female partners, contorting their bodies in forms resembling flowers, ballerinas, and smiling faces^18^. Female *Serromyia femorata* midges pierce and suck conspecific males dry during mating, breaking off his genitalia inside her, thereby supplying the female with both nutrition and sperm^19^. Although an astonishing diversity of sexually dimorphic behaviors exists across species, most insights into the genetic and neural basis of sex-specific behaviors have come from a limited set of model organisms^20^. Which genes control sexual dimorphism in specialist species that have evolved novel behaviors? Do conserved genes control sexual dimorphism in species-specific behaviors, or do novel genes evolve to control new behaviors?

Many advances in understanding the genetics of sexually dimorphic behaviors have come from the study of *Drosophila melanogaster* fly courtship, where a male fly orients towards, taps, and follows a female fly, extending a wing to produce a courtship song before tasting, mounting, and copulating with her^12^. Courtship comprises behavioral modules, which are simple discrete behaviors that must be combined to perform a complex behavior and are elicited by different sensory modalities and subsets of *fruitless-expressing* neurons^15^,^21-24^. Courtship modules include orienting, which is driven by visual information^25^ and persistent following and singing, which are triggered by chemical cues on a female fly^15^ and guided by vision^25,26^. Sex-specific splicing of the *fruitless* gene controls several aspects of courtship behavior. Male flies mutant for *fruitless* promiscuously court other males and cannot successfully mate with females^13,27^. Forcing male *fruitless* splicing in females triggers orientation and singing behaviors normally only performed by males^14^. *fruitless* encodes a BTB zinc-finger transcription factor that is thought to control cell identity and connectivity during development^28,29^, as well as the functional properties of neurons in adulthood^30^, although its genomic targets and the molecular mechanism by which it acts remain unclear. *fruitless* has a conserved role controlling courtship in multiple *Drosophila* species^16,17^, and sex-specific *fruitless* splicing is conserved across wasps^10^ and mosquitoes^9,11^, suggesting that *fruitless* may act as a master regulator of sexually dimorphic mating behaviors across insects.

Mosquitoes display striking sexually dimorphic mating and feeding behaviors. Only male mosquitoes initiate mating, and only females drink blood, which they require to develop their eggs. Sexual dimorphism in blood-feeding is one of the only instances of a completely sexually-dimorphic feeding behavior, since male mosquitoes never pierce skin or engorge on blood. While part of this dimorphism is enforced by sex-specific genitalia^31^ or feeding appendages^32^, there is also a dramatic difference in the drive to hunt hosts between males and female mosquitoes^2,33^. To blood-feed, females combine multiple behavioral modules^2^. Female *Aedes aegypti* mosquitoes take flight when exposed to carbon dioxide^2,6^, and are attracted to human olfactory^4,5,7^, thermal, and visual cues^6,34,35^, and integrate at least two of these cues to orient toward and land on human skin. Engorging on blood is triggered by specific sensory cues tasted by the female^1,8^. It is not known which genes have evolved to control this unique sexually dimorphic and mosquito-specific feeding behavior. Here we generate *fruitless* mutant *Aedes aegypti* mosquitoes and show that consistent with observations in *Drosophila, fruitless* is required for male mating behavior. Unexpectedly, *fruitless* mutant male mosquitoes gain the ability to host-seek, specifically driven by an attraction to human odor. Our results demonstrate that sexual dimorphism in a single module of a mosquito-specific behavior is controlled by a conserved gene that we speculate has gained a new function in the course of evolution.

## RESULTS

### *fruitless* is sex-specifically spliced in the mosquito nervous system

We used an arm-next-to-cage assay (Fig. 1a) to monitor attraction of male and female *Aedes aegypti* mosquitoes to a live human arm. Consistent with their sexually dimorphic blood-feeding behavior, only females were strongly attracted to the arm (Fig. 1b-c). What is the genetic basis for this extreme sexual dimorphism? We reasoned that *fruitless*, which is alternatively spliced in a sex-specific manner and promotes male courtship and copulation in *Drosophila melanogaster* flies^13,27^, may play similar roles in controlling sexually dimorphic behaviors in *Aedes aegypti. fruitless* is a complex gene with multiple promoters and multiple alternatively spliced exons. A previous study showed^11^ and we confirmed that transcripts from the upstream neuron-specific (P1) promoter in the *Aedes aegypti fruitless* gene are sex-specifically spliced (Fig. 1d-h). Both male and female transcripts include a short male ‘m’ exon, and female transcripts additionally include a longer female ‘f’ exon with an early stop codon, predicted to yield a truncated Fruitless protein in the female. However, it is unlikely that any Fruitless protein is stably expressed in the female. In *Drosophila*, female fruitless peptides are not detected^36^, and *transformer* is thought to inhibit translation by binding to female *fruitless* transcripts^37^. P1 transcripts of both sexes splice to the first common ‘c1’ exon, but only male transcripts are predicted to encode full-length Fruitless protein with BTB and zinc-finger domains (Fig. 1e). By analyzing previously published tissuespecific RNA-seq data^38^, we verified that of all the *fruitless* exons, only the f exon was sex-specific in *Aedes aegypti* brains (Fig. 1g). Moreover, P1 transcripts were specifically expressed in the brain and the antenna, the major olfactory organ of the mosquito (Fig. 1h), consistent with *fruitless* P1 expression in *Drosophila*^39^.

**Figure 1.**
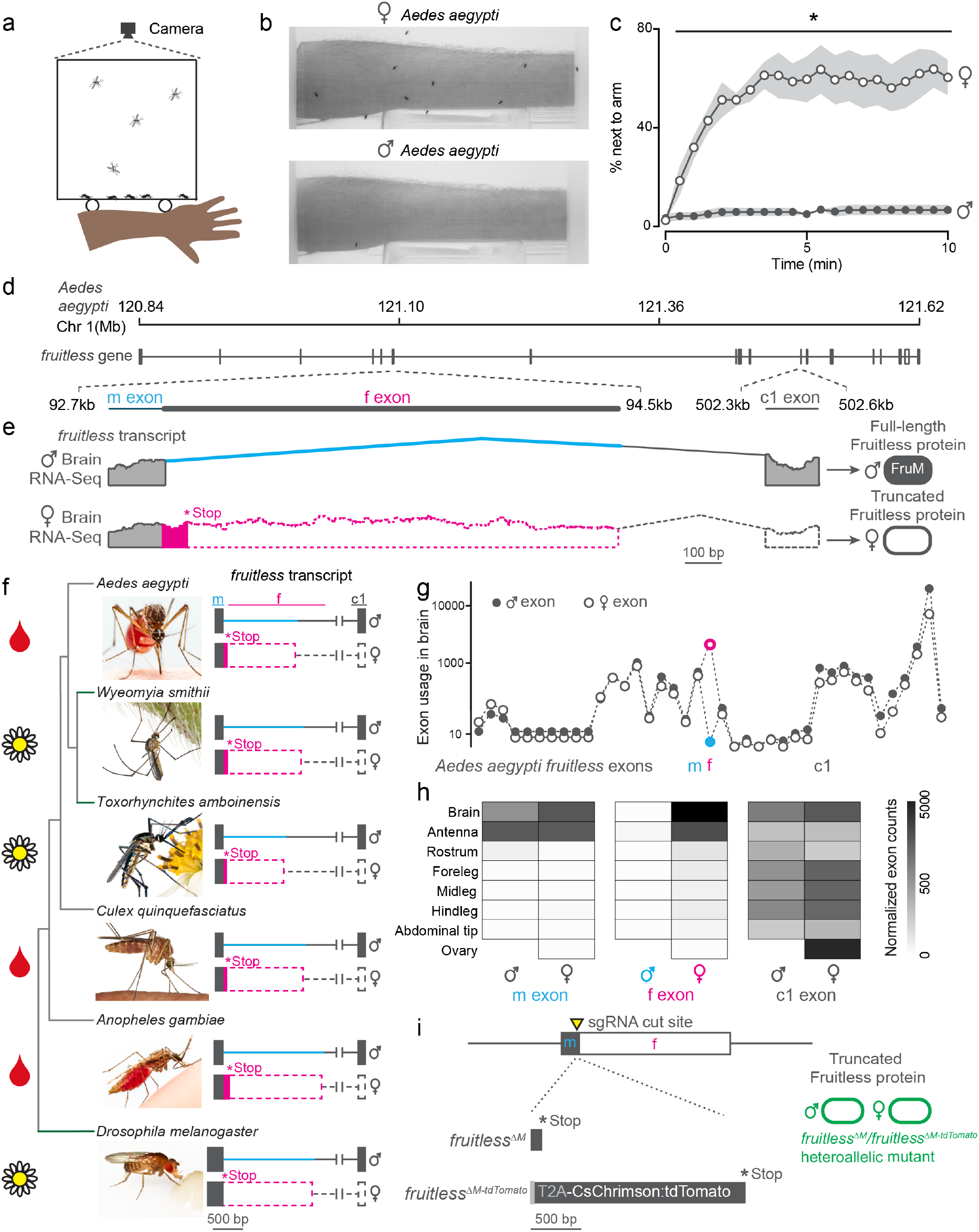
Sex-specific mosquito attraction to humans and *fruitless* splicing. **a, b**, Arm-next-to-cage assay schematic (**a**) and image (**b**) with male (top) and female (bottom) *Aedes aegypti* mosquitoes. **c**, Percent mosquitoes next to arm measured every 30 sec. Data are mean±s.e.m., *n* = 6 trials, *n* = 20 mosquitoes/trial; *p < 0.05, Mann-Whitney test for each time point. **d, e** Schematic of *Aedes aegypti fruitless* genomic locus (**d**) and sex-specific splicing region with RNA-seq read evidence (**e**). **f**, Phylogeny of mosquito species with outgroup *Drosophila melanogaster*, with conserved *fruitless* exon structure inferred from *de novo* transcriptome assembly. In **e, f** coding and non-coding exons are represented by filled and open dashed bars, respectively. *Toxorhynchites rutilus* and *Culex salinarius* images were used to represent *Toxorhynchites amboinensis* or *Culex quinquefasciatus* respectively. See acknowledgments for photo credits. Cartoons indicate blood-feeding (blood drop) and non-blood-feeding (flower) species. **g, h,** *Aedes aegypti fruitless* exon usage based on male and female RNA-seq data (normalized counts) from the indicated tissue plotted for each exon (**g**) and m, f, and c1 exons (**h**) (*n* = 3-4 independent RNA-seq replicates^38^). **i**, Schematic of generation of *fruitless*^Δ*M*^ and *fruitless*^Δ*M-tdTomato*^ mutants.

To ask if *fruitless* splicing was conserved across mosquitoes, we sequenced RNA from male and female brains of five different species and assembled *de novo* transcriptomes for each sex. Three of these species are important arboviral disease vectors because their females blood feed on humans, whereas adults of the two other species only feed on plants^40,41^ (Fig. 1f). We identified orthologues of *fruitless* in each species and found that all had conserved ‘m’ and ‘c1’ exons and distinct ‘f’ exons. *fruitless* was sex-specifically spliced in each of these species with a female-specific ‘f’ exon and early stop codon, predicted to produce a full-length fruitless protein only in males.

We used CRISPR-Cas9 genome editing^42^ to disrupt P1 neural-specific *fruitless* transcripts in *Aedes aegypti* to investigate a possible role of *fruitless* in sexually dimorphic mosquito behaviors. We generated two alleles, *fruitless*^Δ*M*^, which introduces a frameshift that is predicted to produce a truncated protein in males, and *fruitless*^Δ*M-tdTomato*^, in which the *fruitless* gene is disrupted by a knocked-in CsChrimson:tdTomato fusion protein (Fig. 1i). In both alleles, the protein is truncated before the downstream BTB and zinc-finger domains. The *fruitless*^Δ*M-tdTomato*^ line allowed us to visualize cells that express the fluorescent tdTomato reporter under the control of the endogenous *fruitless* regulatory elements. To control for independent background mutations, we used the heteroallelic *fruitless*^Δ*M*^/*fruitless*^Δ*M-tdTomato*^ mutant strain in all subsequent behavior assays (Fig. 1i). In this heteroallelic mutant, *fruitless* P1 transcripts are disrupted in both males and females. Since full-length fruitless protein is male-specific, we expected that only *fruitless*^Δ*M*^/*fruitless*^Δ*M-tdTomato*^ male mosquitoes would display altered behavioral phenotypes.

### Sexually dimorphic expression of *fruitless* in the mosquito brain and antenna

In *Drosophila melanogaster*, P1 *fruitless* transcripts are expressed in several thousand cells comprising about ~2% of the neurons in the adult brain^39^. To examine the distribution of cells expressing *fruitless* in male and female *Aedes aegypti* mosquitoes, we carried out whole mount brain staining to reveal the tdTomato marker expressed from the *fruitless* locus (Fig. 2a-d). *fruitless>tdTomato* is expressed in a large number of cells in both male and female brains (Fig. 2a-d, Fig. S1a-c), as well as in the ventral nerve cord (Fig. S1d-e). *fruitless>tdTomato*-expressing cells innervate multiple regions of the mosquito brain, including the suboesophageal zone, the lateral protocerebral complex, and the lateral horn. These areas have been implicated in feeding^8^, mating^17^, and innate olfactory behaviors^43^ respectively, and also receive projections from *fruitless*-expressing neurons in *Drosophila*^17,39^. The projections of *fruitless>tdTomato* neurons appear to be sexually dimorphic, with denser innervation in the female suboesophageal zone and the male lateral protocerebral complex (Fig. 2b-c, Fig. S1a-b). We did not detect any gross anatomical differences between heterozygous and heteroallelic *fruitless* mutant male brains or the pattern of *fruitless>tdTomato* expression (Fig. 2c-d, Fig. S1b-c). We cannot exclude the possibility that there are subtle differences that can only be observed with sparse reporter expression in subsets of cells.

**Figure 2.**
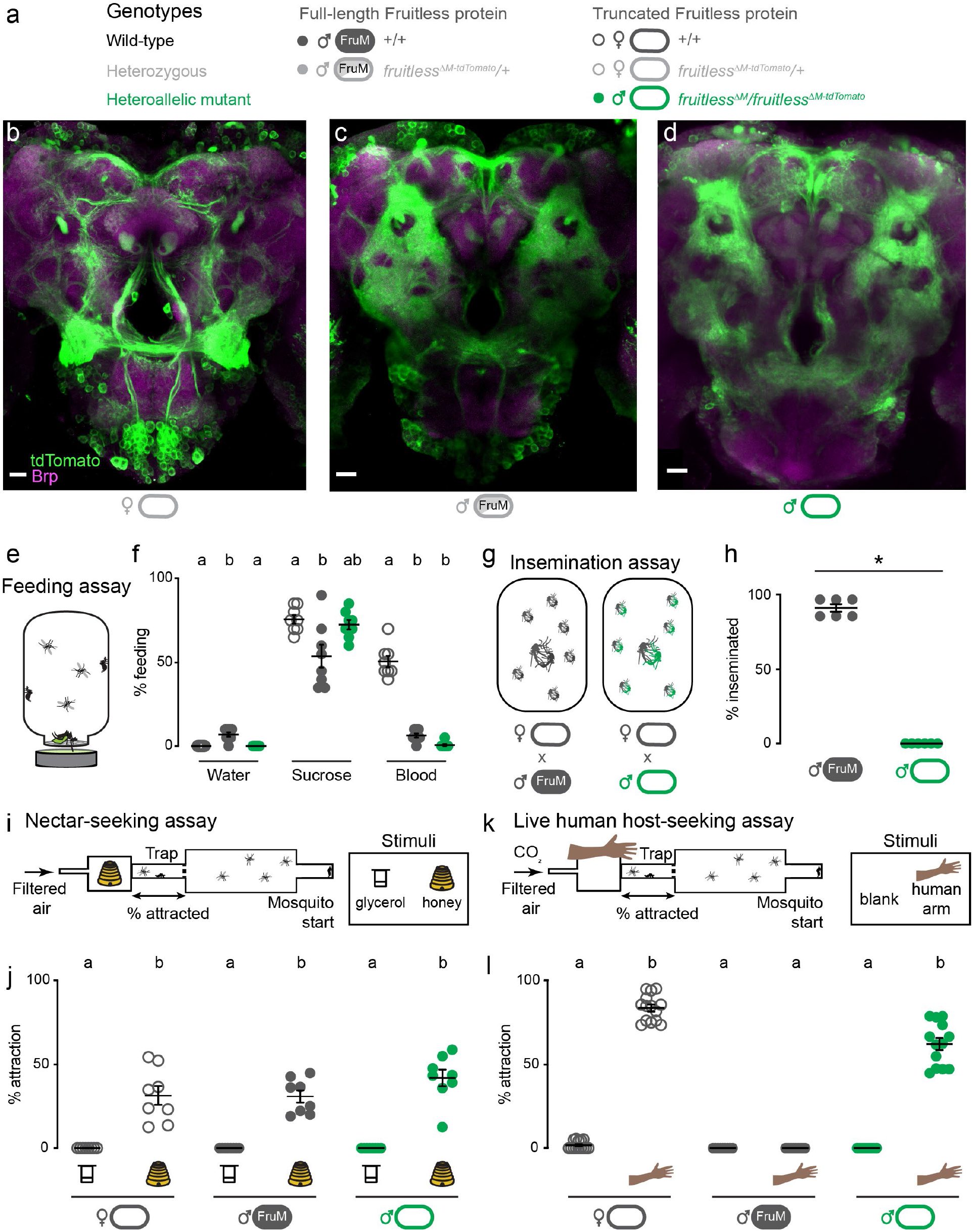
Male *fruitless* mutant mosquitoes gain attraction to a live human host. **a**, Genotypes grouped by effect on Fruitless protein. **b-d**, Confocal images of brains of the indicated genotypes showing *fruitless>tdTomato* (green) and Brp (magenta) expression. Scale bars, 20 μm. **e**, Feeding assay schematic. **f**, Feeding on indicated meal (*n* = 8 trials/meal; *n* = 20 mosquitoes/trial). **g**, Insemination assay schematic. **h**, Insemination of wild-type females by males of the indicated genotype (*n* = 6 trials/male genotype, *n* = 20 females/trial; *p = 0.0022, Mann-Whitney test). **i, k**, Quattroport assay schematic for nectar-seeking (**i**) and live human host seeking (**k**). **j, l** Percent of attracted animals (*n* = 8-14 trials per group, *n* = 17-28 mosquitoes/trial). Data in **f, h, j, l** are mean±s.e.m. In **f, j, l**, data labeled with different letters are significantly different from each other (Kruskal-Wallis test with Dunn’s multiple comparisons, p < 0.05).

We also examined *fruitless* expression in the periphery. Odors are sensed by olfactory sensory neurons in the mosquito antenna, and each type of neuron projects to a single glomerulus in the antennal lobe of the mosquito brain (Fig. S2a). We found that, as is the case in *Drosophila*^39^, *fruitless>tdTomato* is expressed in olfactory sensory neurons in the antenna of both male and female mosquitoes, and that some of these neurons co-express the olfactory receptor coreceptor Orco (Fig. S2b-e). *fruitless>tdTomato* labels a subset of glomeruli in the antennal lobe, with females having about twice as many positive glomeruli compared to males of either genotype (Fig. S2f-l). There was no difference in the number of *fruitless>tdTomato*-labeled glomeruli between wild-type and *fruitless* mutant males (Fig. S2f), suggesting that *fruitless* does not control sexual dimorphism in the number of glomeruli labeled by *fruitless>tdTomato*.

### *fruitless* mutant males gain attraction to live human hosts

Given the broad neural expression and sexual dimorphism in *fruitless* circuits, we asked if *fruitless* mutant males showed any defects in sexually dimorphic feeding and mating behaviors. Since only female mosquitoes have the anatomical capacity to pierce skin and artificial membranes^3,8^, we developed a feeding assay in which both females and males are able to feed from warmed liquids through a net without having to pierce a membrane to access the meal (Fig. 2e). Both wild-type males and females reliably fed on sucrose and did not feed on water. Only wild-type females fed on blood. Even when warm blood was offered and available to males for ready feeding, they still did not find it appetizing (Fig. 2f). *fruitless* mutant males fed similarly to their wild-type male counterparts on all meals, suggesting that this behavioral preference is not under the control of *fruitless* in males (Fig. 2f).

Because *fruitless* plays a key role in male courtship and mating in *Drosophila*, we asked if it is similarly required in *Aedes aegypti*. Since mosquitoes show extremely rapid in-flight mating behavior that is difficult to observe or quantify^44^, we used previously developed insemination assays^45,46^ to quantify the ability of males to successfully mate (Fig. 2g). We found that *fruitless* mutant males were unable to inseminate wild-type females (Fig. 2h). This mating failure is consistent with the established role of *fruitless* in *Drosophila* male sexual behavior^13,14^.

We then turned to innate olfactory behaviors that govern the search for nectar, which is used as a source for metabolic energy by both males and females, and blood, which is required only by females for egg production. Consistent with the use of these meals, nectar-seeking behavior is not sexually dimorphic, but human host-seeking behavior is strictly female specific. To measure these behaviors, we adapted the Uniport olfactometer^47^, which is only able to test one stimulus at a time, and developed the Quattroport, an olfactometer that tests attraction to four separate stimuli in parallel (Fig. 2i,k). To model nectar-seeking behavior, we used honey as a floral odor and glycerol as a control odor as previously described^7^. There was no difference in nectar-seeking between wild-type females, males, and *fruitless* mutant males (Fig. 2j).

We next used the Quattroport with a live human host as a stimulus. As expected, wild-type females robustly and reliably entered traps in response to a live human forearm.

In contrast, zero wild-type males entered the trap, consistent with our observations in the arm-next-to-cage assay (Fig. 1a-c). If *fruitless* function in *Aedes aegypti* were limited to mating and courtship as it is in *Drosophila*, we would expect *fruitless* mutant males to show no interest in a live human host. Unexpectedly, *fruitless* mutant males were as attracted to a live human host as wild-type females (Fig. 2l). This indicates that fruitless males have gained the ability to hostseek, displaying the signature sexually dimorphic behavior of the female mosquito.

### Olfactory and not heat cues attract *fruitless* mutant males to hosts

A live human arm gives off multiple sensory cues that are known to attract female mosquitoes, the most salient of which are body odor and heat. *fruitless* mutant males might be attracted by heat alone or only the human odor, or to the simultaneous presentation of both cues. To disentangle the contribution of these complex sensory cues to the phenotype we observed, we tested the response of *fruitless* mutant males to each cue in isolation. We first used a heatseeking assay^6,48^ to present heat to mosquitoes in the absence of human odor (Fig. 3a-b). Neither *fruitless* mutant nor wild-type males were attracted to the heat cue at any temperature (Fig. 3c). In contrast, wild-type females showed typical heat-seeking behavior that peaked near human skin temperature (Fig. 3c).

**Figure 3.**
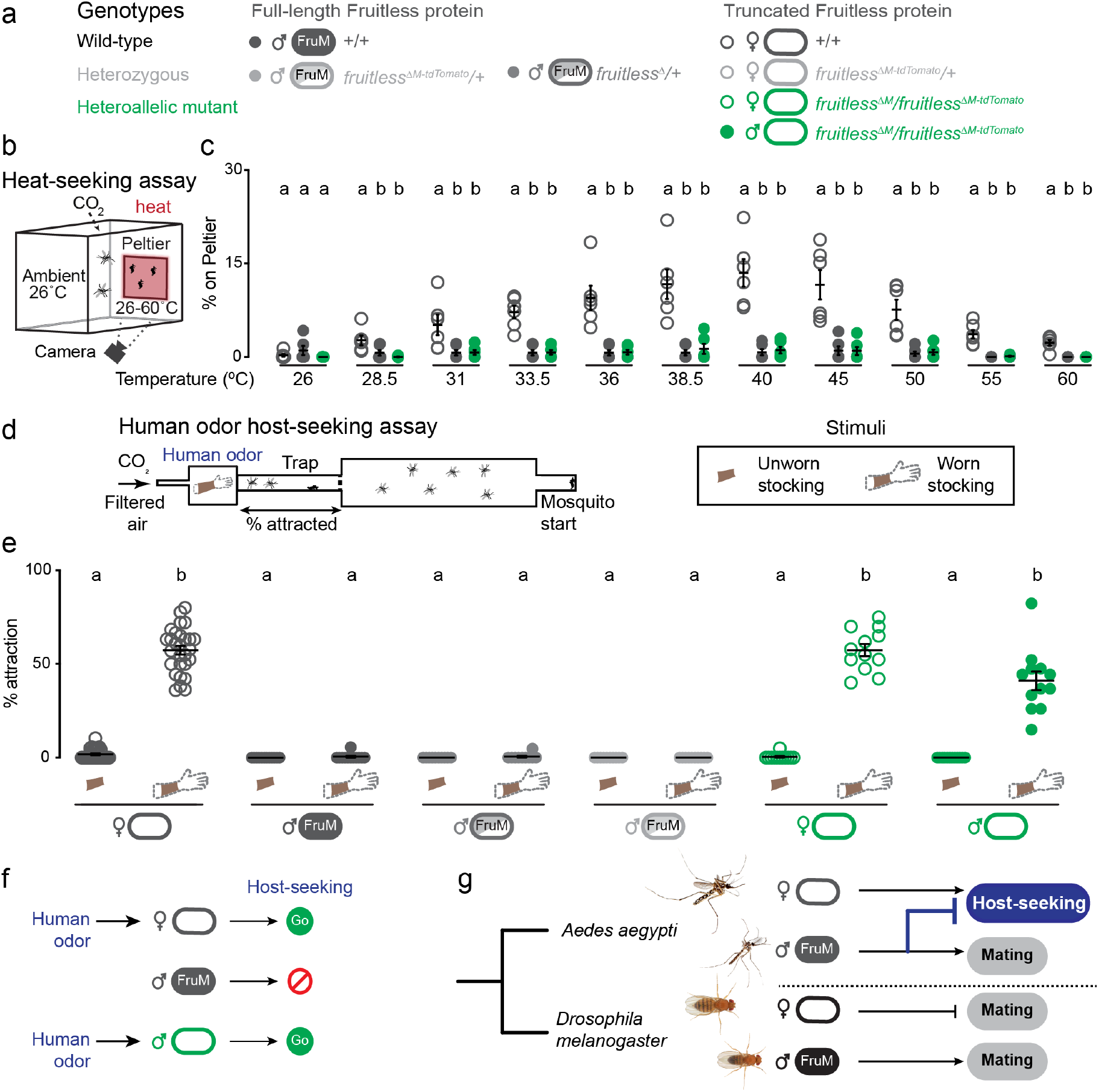
Olfactory cues selectively drive male *fruitless* mutant attraction to humans. **a**, Genotypes grouped by effect on Fruitless protein. **b**, Heat-seeking assay schematic. **c**, Percent of animals on Peltier. Data are mean±s.e.m., *n* = 6 trials/temperature, *n* = 50 mosquitoes/trial. Data labeled with different letters are significantly different from each other, within each temperature. **d**, Schematic of human odor host-seeking assay (left) and stimuli (right). **e**, Percent of attracted animals. Data are mean±s.e.m., *n* = 8-14 trials/group, *n* = 17-28 mosquitoes/trial. **f-g**, Summary of results and model of gain of *fruitless* function in *Aedes aegypti*. Photo credit: *Aedes aegypti* (Alex Wild); *Drosophila melanogaster* (Nicolas Gompel). In **c, e**, data labeled with different letters are significantly different from each other (Kruskal-Wallis test with Dunn’s multiple comparisons, p < 0.05).

To ask if *fruitless* mutant males are attracted to human host odor alone, we collected human scent on nylon stockings and presented this stimulus in the Quattroport (Fig. 3d) to both male and female mosquitoes (Fig. 3a). Whereas wildtype males and heterozygous *fruitless* mutant males showed no response to human odor, wild-type females and *fruitless*^Δ*M*^/*fruitless*^Δ*M-tdTomato*^ females were strongly attracted to human odor (Fig. 3e). Normal host-seeking in *fruitless* mutant females is expected since full-length fruitless protein is translated only in males. Remarkably, heteroallelic *fruitless* mutant males were strongly attracted to human scent, at levels comparable to wild-type females (Fig. 3e). These results demonstrate that *fruitless* mutant males have gained a specific attraction to human odor, which drives them to host-seek.

## DISCUSSION

Only female *Aedes aegypti* mosquitoes host-seek, and we have shown that mutating *fruitless* reveals an attraction to human odor in the male mosquito (Fig. 3f). Previously, *fruitless* was shown to be required for male mating behavior in both *Drosophila*^14^ and *Bombyx* silkmoths^49^. Our work demonstrates that in mosquitoes, *fruitless* has acquired a novel role in inhibiting female host-seeking behavior in the male (Fig. 3g). We cannot exclude the possibility that *fruitless* had a broader ancestral role in repressing male-specific host-seeking or feeding but consider this extremely unlikely given the rarity of sexually dimorphic feeding behaviors relative to sexually dimorphic mating behaviors.

Our results suggest that the neural circuits that promote female attraction to human scent are latent in males and suppressed by expression of *fruitless* either during development, or during adulthood. Since males are able to host-seek in the absence of *fruitless*, the sex of the mosquito does not intrinsically regulate the development and function of brain circuits controlling host-seeking behavior. The concept that a latent sex-specific behavior can be revealed by knocking out a single gene was also demonstrated in the mouse (*Mus musculus*). Only male mice court and initiate sexual contact with females and knocking out the *Trpc2* gene causes female mice to display these male-specific behaviors^50^.

Host-seeking is the first step in a complex sequence of behaviors that lead to blood-feeding. After detecting and flying toward a human host, the female mosquito must land on the human, pierce the skin, and ultimately engorge on blood^2^. We have shown that *fruitless* has evolved to control sexual dimorphism in one module of this specialized behavior, the ability to host-seek. Females integrate multiple sensory cues to identify and approach human hosts, and we show that *fruitless* controls the response to just one of those cues, human odor. Sexual dimorphism in thermosensation, or in subsequent feeding behaviors does not appear to be controlled by *fruitless* in the male mosquito, since neither wild-type nor mutant *fruitless* males will drink warm blood (Fig. 2e,f). The modular genetic organization of mosquito behavior sparks intriguing parallels to other complex sexually dimorphic behaviors like mouse parenting^51^. To be effective parents, female mice must build nests, and then retrieve, groom, and nurse their pups. In the deer mouse *Peromyscus*, the conserved peptide vasopressin has evolved to control nest building^52^. In both *Peromyscus* and *Aedes*, a conserved gene has gained control over a single aspect of a complex behavior. It has been hypothesized that certain classes of genes like neuromodulators and transcription factors are more likely to underlie phenotypic differences between species^53-55^, and our study demonstrates that this is true even for an entirely novel behavior.

Where in the nervous system is *fruitless* required to suppress host seeking in male mosquitoes? *fruitless* might function in the antenna to modulate the detection of human odor in male mosquitoes, perhaps by tuning olfactory sensory neuron function to human cues. However, we do not detect any difference in the number or position of antennal lobe glomeruli between wild-type and *fruitless* mutant males (Fig. S2f), indicating that there is not likely to be a simple peripheral change in the mutants. Alternatively, we favor the model where both males and females detect human odor, and *fruitless* functions in the central brain to reroute these signals to drive different motor outputs, as has been demonstrated with the sexually dimorphic response to *Drosophila* pheromones^22,24,43^. To distinguish between these two models, we require a *fruitless* driver line, and/or the ability to subset expression to label and drive reporters or rescue *fruitless* expression in sparse populations of neurons. Despite significant effort, we were unable to generate a viable *fruitless* driver line (Table S1). Advances in mosquito genetic tools, such as the successful implementation of orthogonal transcriptional activator reagents, combined with sparse labeling approaches will be required to gain mechanistic insight into *fruitless* function within mosquito host-seeking circuits. We note that these advances were not trivial in *Drosophila melanogaster*, requiring efforts from multiple laboratories over the past decade^22,24,43^, and expect that the generation of these tools and the subsequent characterization of the circuit will be significantly more challenging in the mosquito, a non-model organism.

Our work demonstrates that *fruitless* has evolved the novel function of enforcing female-specific host-seeking while maintaining its presumably ancestral male mating function. How might *fruitless* have evolved to control host-seeking? One possibility is that non-sex-specific host-seeking neural circuits first emerged in the ancestral mosquito, and then secondarily began to express *fruitless* to suppress the development or adult function of host-seeking circuits specifically in males. Another possibility is that female mosquitoes duplicated and co-opted the ancestral *fruitless-*expressing mating neural circuits, retuned the inputs and outputs into this circuit to drive host-seeking. In this scenario, *fruitless* function would have switched from promoting mating to inhibiting host-seeking in males. We speculate that both of these possibilities allow for *fruitless* to control both sex-specific host-seeking and mating behaviors, and identification and molecular profiling of the *fruitless* cells controlling host-seeking and mating will help distinguish between these models. Our work highlights *fruitless* as a potential means to investigate the circuit basis of *Aedes aegypti* host seeking, a behavior that is responsible for infecting millions of people with life-threatening pathogens.

## SUPPLEMENTAL INFORMATION

All raw data are provided in Data File 1. Plasmids are available from Addgene (#141099, #141100) and RNA-seq reads are at NCBI SRA Bioproject PRJNA612100. Details of Quattroport fabrication and operation are available at Github: https://github.com/VosshallLab/Basrur_Vosshall2020

## ACKNOWLEDGMENTS

We thank Richard Benton, Josie Clowney, Vanessa Ruta, Nilay Yapici, and members of the Vosshall Lab for comments on the manuscript; Gloria Gordon and Libby Mejia for expert mosquito rearing; Joel Butterwick and Vanessa Ruta for sharing unpublished anti-*Apocrypta bakeri* Orco monoclonal antibodies; the colleagues who shared mosquito species with us [William Bradshaw and Christina Holzapfel (*Wyeomyia smithii*); Larry Zwiebel and Jason Pitts (*Toxorhynchites amboinensis*); Flaminia Catteruccia (*Anopheles gambiae*)] and those who shared insect photos with us [Alex Wild (*Aedes aegypti, Anopheles gambiae*), Lawrence Reeves of Florida Medical Entomology Laboratory (*Wyeomyia smithii, Toxorhynchites rutilus, Culex salinarius*), Francisco Romero, Veronica Corrales-Carvajal, Carlos Ribeiro, and Nicolas Gompel (*Drosophila melanogaster*)]; Erich Jarvis, Vanessa Ruta, and Li Zhao for advice and discussion; Ben Matthews and Zach Gilbert for assistance in designing and testing of sgRNAs and advice on the CRISPR protocol; Ben Matthews for advice on bioinformatics; Rob A Harrell II at the Insect Transgenesis Facility at the University of Maryland for CRISPR-Cas9 injections; Jim Petrillo and Kunal Shah at the Rockefeller Precision Instrument Technology center for help in development and construction of the Quattroport assay; Meg Younger for advice on brain dissections and antibody staining; Ella Jacobs for training in spermathecae dissections; Christina Pyrgaki, Carlos Rico, and Alison North at the Rockefeller Bio-Imaging Resource Center for assistance with confocal imaging; Connie Zhao at the Rockefeller Genomics Resource Center for advice on RNA-seq library preparation and sequencing; Román Corfas for assistance with the heat-seeking assay.

This work was supported in part by grant # UL1 TR000043 from the National Center for Advancing Translational Sciences (NCATS, National Institutes of Health (NIH) Clinical and Translational Science Award (CTSA) program. Harvey L. Karp Discovery Award, Japan Society for Promotion of Science Overseas Research Fellowship to T.M. Harvey L. Karp Discovery Award to M.E.D., and Helen Hay Whitney Foundation Fellowship to M.E.D. This publication was supported in part by the National Center for Advancing Translational Sciences, National Institutes of Health, through Rockefeller University, Grant # UL1 TR001866 to M.E.D. and NIH NIDCD grant F30DC017658 to M.H. M.H. is supported by a Medical Scientist Training Program grant from the National Institute of General Medical Sciences of the NIH under award number T32GM007739 to the Weill Cornell/Rockefeller/Sloan Kettering Tri-Institutional MD-PhD Program. L.B.V. is an investigator of the Howard Hughes Medical Institute.

## AUTHOR CONTRIBUTIONS

N.S.B carried out all experiments with the following exceptions: M.E.D developed and optimized the Quattroport assay, in Fig. 2i-l and Fig3d,e, Y.N.T. developed the feeding assay in Fig. 2e,f, T.M. carried out experiments in Fig. 3b,c, M.H. developed and optimized the protocol for whole mount antibody staining in Fig. S2b-e, R.K.v.H. carried out experiments in Fig. S3d-i. N.S.B and L.B.V. together conceived the study, designed the figures, and wrote the paper with input from all authors.

## MATERIALS AND METHODS

### Human and animal ethics statement

Blood-feeding procedures and behavioral experiments with live hosts were approved and monitored by The Rockefeller University Institutional Animal Care and Use Committee (IACUC protocol 17018) and Institutional Review Board (IRB protocol LV-0652), respectively. Human subjects gave their written informed consent to participate.

### Mosquito rearing and maintenance

*Aedes aegypti* wild-type laboratory strains (Liverpool-IB12) were maintained and reared at 25 - 28°C, 70-80% relative humidity with a photoperiod of 14 hr light: 10 hr dark (lights on at 7 a.m.) as previously described^7^. All behavioral assays were performed at these conditions of temperature and humidity. Adult females were blood-fed on mice for stock maintenance and on human subjects for initial stages of mutant generation. *Anopheles gambiae* (G3 strain), *Wyeomyia smithii* (PB strain), *Toxorhynchites amboinensis*, and *Culex quinquefasciatus* (JHB strain) were reared in similar conditions, following previously described protocols for each species^40,41,56^. Adult mosquitoes of each species were provided constant access to 10% sucrose.

### RNA-sequencing

7 to 14 day-old mosquitoes of each species were cold-anesthetized and kept on ice for up to 1hr or until dissections were complete. Brains were dissected in ice-cold RNase-free phosphate-buffered saline (PBS) (Invitrogen AM9625) on ice, moved into a microfuge tube with forceps, and immediately snap frozen in a cold block (Simport S700-14) chilled to −80°C on dry ice. Each sample group was dissected in parallel to avoid artefacts and batch effects, and five brains were used per sample. Dissected tissue was stored at −80°C until RNA extraction. RNA extraction was performed using the PicoPure Kit (ThermoFisher #KIT0204) following the manufacturer’s instructions, including DNase treatment. Samples were run on a Bioanalyzer RNA Pico Chip (Agilent #5067-1513) to determine RNA quantity and quality. Libraries were prepared using the Illumina TruSeq Stranded mRNA kit #20020594, following manufacturer’s instructions. Library quantity and quality were evaluated using High Sensitivity DNA ScreenTape Analysis (Agilent #5067-5585) prior to pooling. Bar-coded samples from all non-*Aedes* tissues were pooled in an equal ratio before distributing the pool across 2 sequencing lanes. Sequencing was performed at The Rockefeller University Genomics Resource Center on a NextSeq 500 sequencer (Illumina). All reads were 2 x 150 bp. Data were demultiplexed and delivered as fastq files for each library. Sequencing reads have been deposited at the NCBI Sequence Read Archive (SRA) under BioProject PRJNA612100.

### *fruitless* splicing analysis

Reads from individual *Aedes* libraries were mapped to the AaegL5 genome^57^ using STAR version 2.5.2a with default settings^58^. Raw counts were used for differential splicing analysis in *Aedes aegypti* using DEXSeq version 1.32.0 (ref. ^59^) as per author instructions. For the other mosquito species without genomes or incomplete genome annotations, we assembled sex-specific *de novo* transcriptomes using Trinity version 2013-03-25 with default settings^60^. We then searched for *fruitless* orthologues in each species using BLAST 2.6.0 (ref. ^61^), aligned hits to *Aedes aegypti fruitless* P1 transcripts using MacVector version 15.0.3, and picked the best match for each exon, species, and sex. *fruitless* exon sequences are found in Data File 1.

### *fruitless*^Δ*M*^ and *fruitless*^Δ*M-tdTomato*^ strain generation

The *fruitless* gene was targeted using CRISPR-Cas9 methods as previously described^42^. Gene-targeting reagents were injected into wild-type Liverpool-IB12 embryos at the Insect Transformation Facility at the University of Maryland Institute for Bioscience & Biotechnology Research. For each line, either 2000 or 1000 embryos were injected with 600 ng/μL plasmid, 300 ng/μL Cas9 protein (PNABio CP01-200), and 40 ng/μL sgRNA. Proper integration was confirmed in each strain using polymerase chain reaction (PCR) and sequencing. Animals were then back-crossed to wild-type Liverpool-IB12 for at least four generations. All homology arms for homology-directed integration were isolated by PCR using Liverpool-IB12 genomic DNA. sgRNA DNA template was prepared by annealing oligonucleotides as previously described^42^. *In vitro* transcription of sgRNA template was performed using HiScribe Quick T7 kit (New England Biolabs #E2050S) following the manufacturer’s directions and incubating for 4 hr at 37°C. Following transcription and DNAse treatment for 15 min at 37°C, sgRNA was purified using Ampure RNAse-free SPRI beads (Beckman-Coulter #A63987) and eluted in Ultrapure water (Invitrogen #10977–015). For all plasmids, fragments were generated by PCR from the indicated template with the indicated primers (Data File 1) and assembled using NEBuilder HiFi DNA Assembly (NEB E5520S). Plasmids were transformed into NEB competent cells (NEB C2987I), purified with the NucleoBond Xtra Midi Endotoxin Free kit (Clontech 740420.50), and sequence verified. The *fruitless*^Δ*M*^ mutant was generated in the course of attempting to generate a *fruitless* QF2 knock-in mutant (Table S1) (see below). One of the families had viable 3xP3-dsRed positive offspring and an out-of-frame QF2 insertion, which was predicted to produce a truncated fruitless protein in males. This was the *fruitless*^Δ*M*^ mutant allele we used in the study. The *fruitless*^Δ*M-tdTomato*^ knock-in/knock-out strain was generated by inserting a cassette containing T2A followed by CsChrimson fused to the fluorescent protein tdTomato and the 3xP3-EYFP strain marker. We obtained 2 independent viable lines and selected one for use in this study. We used the CsChrimson:tdTomato protein expressed from the *fruitless* locus in *fruitless*^Δ*M-tdTomato*^ animals as a marker for *fruitless* expression in these studies. CsChrimson is a red light-activated cation channel and we originally generated this animal with the intention of optogenetically manipulating behavior. However, CsChrimson-tdTomato was not detectable in the brain without antibody amplification from antibody staining, suggesting that expression is too low for optogenetic stimulation.

### Attempted generation of *fruitless* QF2/QF2w knock-in mutants

We attempted to generate *fruitless* P1-specific driver lines by knocking in a cassette containing the ribosomal-skipping peptide T2A followed by the transcriptional activator QF2 (ref. ^62^), with 3xP3-dsRed as an insertion marker as previously described^63^. In this knock-in/knock-out strain we aimed to disrupt the *fruitless* gene as well as generate a driver line that would allow us to label and manipulate *fruitless*-expressing neurons. We recovered 7 independent 3xP3-dsRed positive G1 families. However, all females with one copy of the correct integration did not blood-feed after many attempts using multiple different human hosts. Males with one copy of this insertion did not mate with wild-type females. Since blood-meals are required for *Aedes aegypti* egg-development, this line could not be maintained. We next tried to knock-in the weaker QF2w transcriptional activator, and recovered 6 independent families, all of which showed the same blood-feeding and mating defects (Table S1). We speculate that toxicity of QF2 or QF2w may affect the function or viability of *fruitless*-expressing neurons, leading to the behavioral defects we observed. The cause of Q-system toxicity, even attenuated from Q to QF2 to QF2w is unknown^62^. We speculate that this toxicity is unrelated to the *fruitless* locus, because *fruitless*^Δ*M*/+^ animals had no phenotype as heterozygotes, unlike *fruitless*^Δ*QF2*/+^ and *fruitless*^Δ*QF2w*/+^ animals.

### Attempted generation of sex-switched *fruitless* mutants

To ask if fruitless protein was sufficient to inhibit host-seeking behavior in females, we attempted to force females to express male fruitless protein by deleting the female exon of *fruitless* P1 transcripts and forcing male *fruitless* splicing in female brains (Table S1). For the *fruitless*^Δ*F*^ line, embryos were injected with 300 ng/μL Cas9 protein, 125 ng/μL oligonucleotide with template repairing the splice site, 40 ng/μL each of two sgRNAs targeting the beginning and end of the female-specific exon. We recovered multiple G1 animals with the correct integration, as verified by PCR and sequencing. Male *fruitless* splicing in *fruitless*^Δ^ females was verified with reverse-transcription PCR (data not shown). G2 *fruitless*^Δ*F*^ females did not fully blood-feed or lay eggs even though they were successfully inseminated by wild-type males (Figure S3d-i). It was therefore impossible to maintain these lines. We do not know if the bloodfeeding defect was due to a failure to respond to the host or some other behavioral or anatomical defect. Since *fruitless* is tightly linked to the male-determining locus, it was not an option to maintain this targeted allele in males. Integrations on the male chromosome would yield ~1/500 females with the recombinant allele, and integrations on the female chromosome yield inviable females and rare recombinant males. In either scenario, the *fruitless*^Δ*F*^ insertion is unmarked and would need to be followed by PCR genotyping. We also attempted to generate a line where we both deleted the female *fruitless* exon and knocked-in an intronic 3xP3 fluorescent marker, which would allow us to maintain this line in males and use the marker to select rare recombinants for behavioral analysis. However, females with this integration did not have any behavioral phenotypes, suggesting that the intronic 3xP3 marker interfered with regular *fruitless* splicing in both males and females. These difficulties precluded any further investigation of the phenotype of expressing full-length fruitless protein in females.

### Antibody staining – brain whole mounts

Dissection of adult brains and immunostaining was carried out as previously described^8,63^. 6 to 14 day-old mosquitoes were anesthetized on ice and decapitated. Heads were fixed in 4% paraformaldehyde (Electron Microscopy Sciences 15713-S), 1X Ca^+2^, Mg^+2^ free PBS (Thermo 14190144), 0.25% Triton X-100 (Sigma 93443), and nutated for 3 hr at 4°C. Brains were then dissected and placed in cell-strainer caps (Falcon #352235) in a 24 well-plate. All subsequent steps were performed on a low-speed orbital shaker. Brains were washed for 15 min at room temperature in 1x PBS with 0.25% Triton X-100 (0.25% PBT) at least 6 times. Brains were permeabilized with 4% Triton X-100 with 2% normal goat serum (Jackson ImmunoResearch #005-000-121) in PBS at 4°C for 2 days. Brains were washed for 15 min at least 6 times with 0.25%PBT at room temperature. Brains were incubated in 0.25%PBT plus 2% normal goat serum with primary antibodies at the following dilutions: rabbit anti-RFP (Rockland 600-401-379) 1:200 and mouse anti-*Drosophila melanogaster* Brp (nc82) 1:5000. The nc82 hybridoma developed by Erich Buchner of Universitätsklinikum Würzburg was obtained from the Developmental Studies Hybridoma Bank, created by the NICHD of the NIH and maintained at The University of Iowa, Department of Biology, Iowa City, IA 52242. Primary antibodies were incubated for 2 nights at 4°C degrees then washed at least 6 times for 15 min with 0.25% PBT at room temperature. Brains were incubated with secondary antibody for 2 nights at 4°C with secondary antibodies at 1:500 and 2% normal goat serum. Secondary antibodies used were goat anti-rabbit Alexa Fluor 555 (Thermo A32732) and goat anti-mouse Alexa Fluor 647 (Thermo A-21235). Brains were then washed for 15 min at least 6 times with 0.25% PBT at room temperature and mounted in Slowfade Diamond (Thermo S36972) using #1.5 coverslips as spacers before confocal imaging.

### Antibody staining – antennal whole mounts

This protocol was adapted from a *Drosophila* embryo staining protocol^64^. 6 to 10 day-old mosquitoes were anesthetized, decapitated, and placed in 1.5 mL 5 U/mL chitinase (Sigma C6137) and 100 U/mL chymotrypsin (Sigma CHY5S) in 119 mM NaCl, 48 mM KCl, 2 mM CaCl2, 2 mM MgCl2, 25mM HEPES buffer on ice. Male heads were incubated for 5 min on a ThermoMixer (Eppendorf 5382000023), and 25 min in a rotating hybridization oven, and female heads were incubated for 10 min on the ThermoMixer and 50 min in rotating oven, all at 37°C. Heads were then rinsed once and fixed in 4% paraformaldehyde, 1X Ca^+2^, Mg^+2^ free PBS, and 0.25% Triton X-100 for 24 hr at room temperature on a rotator. All subsequent 4°C steps used a nutator, and room temperature steps used a rotator. Heads were washed for 30 min at room temperature at least 3 times in 1X PBS with 0.03% Triton X-100 (0.03% PBT). Antennae were then dissected into 0.5 mL microfuge tubes and dehydrated in 80% methanol / 20% DMSO for 1 hr at room temperature. Antennae were washed in 0.03% PBT for 30 min at room temperature, and blocked/permeabilized in 1X PBS, 1% DMSO (Sigma 472301), 5% normal goat serum, 4% Triton X-100 for 24 hr at 4°C. Antennae were washed for 30 min at least 5 times with 0.03% PBT, 1% DMSO at room temperature, and then moved to primary antibody in 1X PBS, 1% DMSO, 5% normal goat serum, 0.03% Triton X-100 for 72 hr at 4°C. Primary antibodies used were mouse anti-*Apocrypta bakeri* Orco monoclonal antibody #15B2 (1:50 dilution, gift of Joel Butterwick and Vanessa Ruta), and rabbit anti-RFP (1:100, Rockland 600-401-379). Orco monoclonal antibody specificity was verified in *Aedes aegypti* by staining *orco* mutant antennae, which showed no staining (data not shown). Antennae were washed for 30 min at least 5 times with 0.03% PBT, 1% DMSO at room temperature, and then washed overnight in the same solution. Antennae were then moved to secondary antibody (1:200) and DAPI (1:10000, Sigma D9542) in 1X PBS, 1% DMSO, 5% normal goat serum, 0.03% Triton X-100 for 72 hr at 4°C. Secondary antibodies used were goat anti-mouse Alexa Fluor 488 (Thermo A-11001) and goat anti-rabbit Alexa Fluor 555 Plus (Thermo A32732). Antennae were washed for 30 min at least 5 times with 0.03% PBT, 1% DMSO at room temperature, and then washed overnight in the same solution. Antennae were rinsed in 1X PBS, rinsed 3 times in Slowfade Diamond (Thermo S36972), and mounted in Slowfade Diamond.

### Confocal image acquisition

Images were acquired with a Zeiss Axio Observer Z1 Inverted LSM 880 NLO laser scanning confocal microscope (Zeiss) with either 25x/0.8 NA (whole brains) or 40x/1.4 NA (antennal lobes, antennae) immersion-corrected objective at a resolution of 1024 x 1024 or 2048 x 2048 (brains) or 3024 x 1024 (antennae) pixels. Confocal images were processed in ImageJ (NIH).

### Arm-next-to-cage assay

This assay was performed as described previously^7^. Briefly, for each trial, 20 adult mosquitoes were sorted under cold anesthesia (4°C) and placed in a cage and allowed to acclimate for 30 min. A human arm was placed 2.5 cm from one side of a standard 28 x 28 x 28 cm cage. Mosquitoes could not directly contact the human arm. A Logitech C920s HD Pro Webcam was positioned to take images of mosquitoes responding to the human arm. Trials ran for 10 min and images were acquired at a rate of 1 frame per sec. To quantify mosquito responses, we manually counted the number of mosquitoes resting on the lower portion of the screen closest to the human arm.

### Feeding assay

Mosquitoes were cold-anesthetized, and 20 mosquitoes were sorted into 250 mL bottles covered with a taut net secured with rubber bands. They were allowed to acclimate for 24 hr with access to water through cotton balls. The following meals were presented: water, 10% sucrose, or sheep’s blood (Hemostat DSB100) supplemented with 1mM ATP (Sigma A6419). Meals were warmed to 45°C before being used in the assay. 10 mL of a given meal was pipetted into the bottle caps, animals were activated with a 30 sec pulse of 4% CO_2_, and bottles were inverted on top of the caps. Mosquitoes were allowed to feed on each meal through the net for 10 min and were then anesthetized at 4°C and scored as fed if any level of feeding was observed, as assessed by visual inspection of the abdomen of the animal.

### Insemination assay

Mosquitoes were separated by sex at the pupal stage and sex was confirmed within 24 hr of eclosion. For each trial, 10 female Liver-pool-IB12 virgin mosquitoes were crossed to 11 virgin male mosquitoes of either Liverpool-IB12 or *fruitless*^Δ*M*^/*fruitless*^Δ*M-tdTomato*^ genotype in a bucket cage for 24 hr, with access to 10% sucrose. Mosquitoes were then anesthetized at 4°C, females separated from males, and female spermathecae were dissected to score for insemination as a sign of successful mating^46^. Control virgin females were dissected in parallel to verify absence of insemination.

### Quattroport olfactometer

Details of Quattroport fabrication and operation are available at https://github.com/VosshallLab/Basrur_Vosshall2020. Briefly, the Quattroport consists of four tubes, each with its own stimulus box, trap, and mosquito start chamber. There are adjustable gates between each chamber. The stimulus was placed upstream of a trap, and mosquitoes are prevented from contacting the stimulus by a mesh barrier. In each trial, four stimuli were run in parallel, with the positions of stimuli randomized and rotated between each trial. Air was filtered and pumped into each box, either in the presence of CO_2_ (for host-seeking assays) or without CO_2_ (honey-seeking assays). For all assays, ~20 mosquitoes were sorted and placed into canisters the day of behavior. Mosquitoes were allowed to acclimate in the assay for 10 min, then exposed to the stimulus for 30 sec, after which gates were opened and animals allowed to fly for 5 min. After this time, gates were closed and mosquitoes were counted to quantify the percent of mosquitoes in the trap. For honey assays, 3 to 7 day-old mosquitoes were fasted for 24 hr before the experiment by replacing 10% sucrose with a water source. CO_2_ was not supplied for honey assays. Either 1 g of leatherwood honey (Tasmanian Honey Company, Tasmania, Australia) or glycerol (Sigma G5516) was applied to a 55 mm diameter Whatman filter paper circle (GE Healthcare, Buckinghamshire, UK) and placed in a Petri dish. For host-seeking assays, mosquitoes were allowed access to sucrose before the experiment. CO_2_ was supplied in the airstream for the duration of the 5 min 30 sec assay in all host-seeking assays (for both human forearm/odor stimuli and blank/unworn nylon controls). For live human host-seeking assays, a human subject placed their forearm on an acrylic box, exposing a 2.5 x 5 cm rectangle of skin to the airstream. For human odor host-seeking assays, the same human subject wore a tan nylon sleeve (L’eggs Women’s Comfortable Everyday Knee Highs Reinforced Toe Panty Hose, modified with scissors to remove the toe area) on their forearm. A second black nylon sleeve was worn on top of the ta n nylon odor sampling sleeve to protect it from external odors. After 6 hr of continuous wear, the black nylon sleeve was discarded, and the tan nylon sleeve was frozen at −20°C. Nylons were used within one week of being worn. On the day of the assay, nylons were thawed for at least 1 hr at room temperature. A 10 x 14 cm piece of the sleeve was presented with the skin-contacting surface facing upward in the stimulus box along with CO_2_. Unworn nylons were similarly frozen, thawed, and cut to serve as negative controls.

### Heat-seeking assay

Experiments were performed as previously described^6,48^. Briefly, 45-50 mosquitoes were fasted for 3 hr before the experiment by replacing 10% sucrose with a water source and were then transferred into a custom-made Plexiglass box (30 x 30 x 30 cm), with carbon-filtered air pumped continuously into the box via a diffusion pad installed on the ceiling of the enclosure. All stimulus periods lasted 3 min and were presented on a single Peltier element (6 x 9 cm, Tellurex) covered with a piece of standard white letter-size printer paper (NMP1120, Navigator) cut to 15 x 17 cm and held taut by a magnetic frame. CO_2_ pulses (20 sec, to >1000 ppm above background levels) were added to the air stream and accompanied all stimulus period onsets. Mosquito landings on the Peltier were monitored by fixed cameras (FFMV-03M2M-CS, Point Grey Research) with images acquired at 1 Hz. Images were analyzed using custom MATLAB scripts to count mosquito landings within a fixed target region. Mosquito occupancy on the Peltier was quantified during seconds 90-180 of each stimulus period.

## QUANTIFICATION AND STATISTICAL ANALYSIS

All statistical analysis was performed using GraphPad Prism Version 8. Data collected as percent of total are shown as mean±s.e.m. Details of statistical methods are reported in the figure legends.

## DATA AND SOFTWARE AVAILABILITY

All raw data are provided in Data File 1. Plasmids are available at Addgene (#141099, #141100). RNA-seq data are available in the Short Read Archive at Genbank (Bioproject: PRJNA612100). Details of Quattroport fabrication and operation are available at Github: https://github.com/VosshallLab/Basrur_Vosshall2020

## SUPPLEMENTARY FIGURES

**Figure S1.**
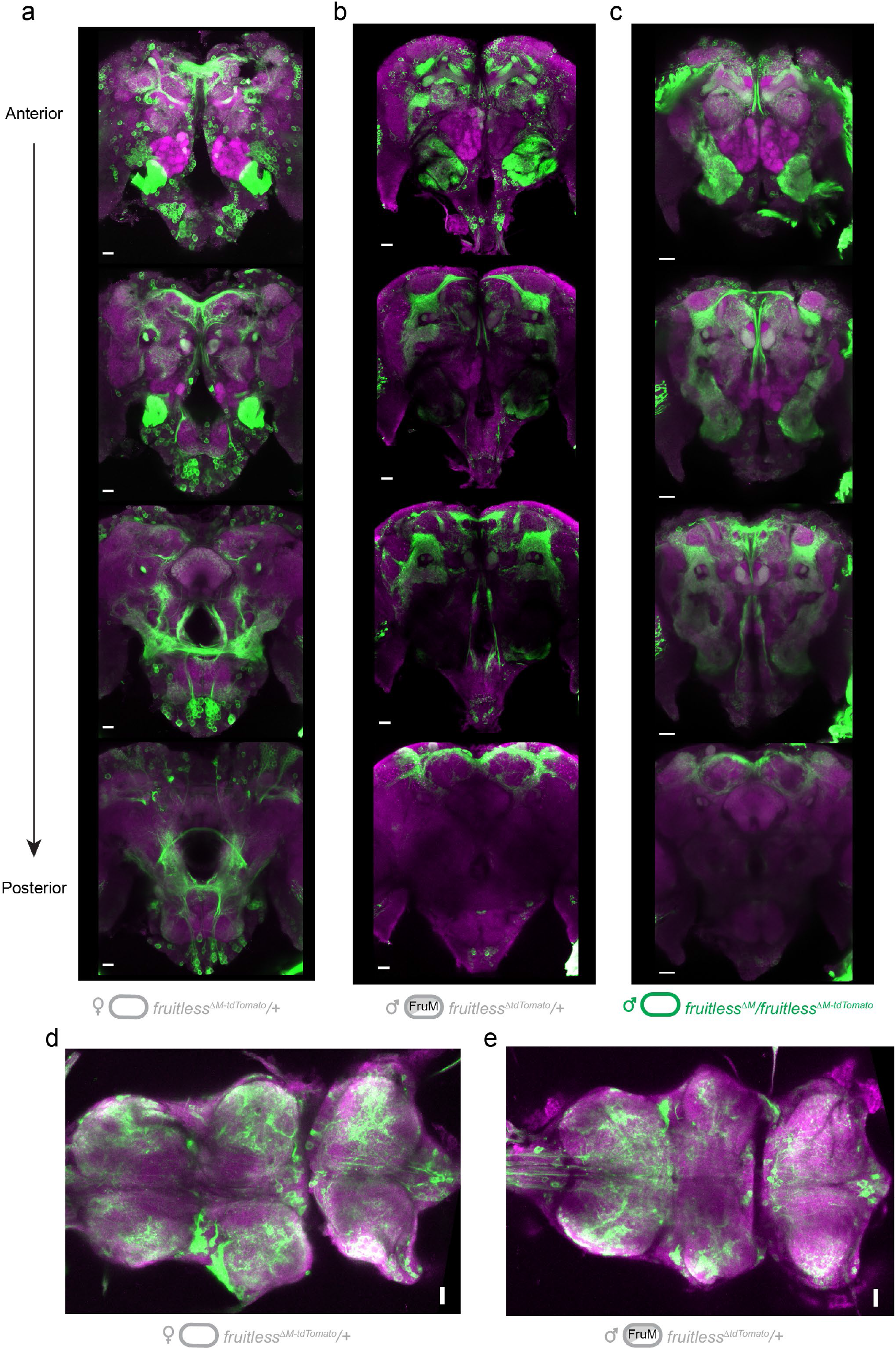
Expression of *fruitless* in the mosquito central nervous system. **a-c**, Confocal images of brains of the indicated genotypes showing *fruitless>tdTomato* (green) and Brp (magenta) expression. Top-to bottom images are optical sections of the same brain, arranged from anterior to posterior. **d-e**, Confocal images of ventral nerve cords of the indicated genotypes showing *fruitless>tdTomato* (green) and Brp (magenta) expression. All scale bars, 20 μm.

**Figure S2.**
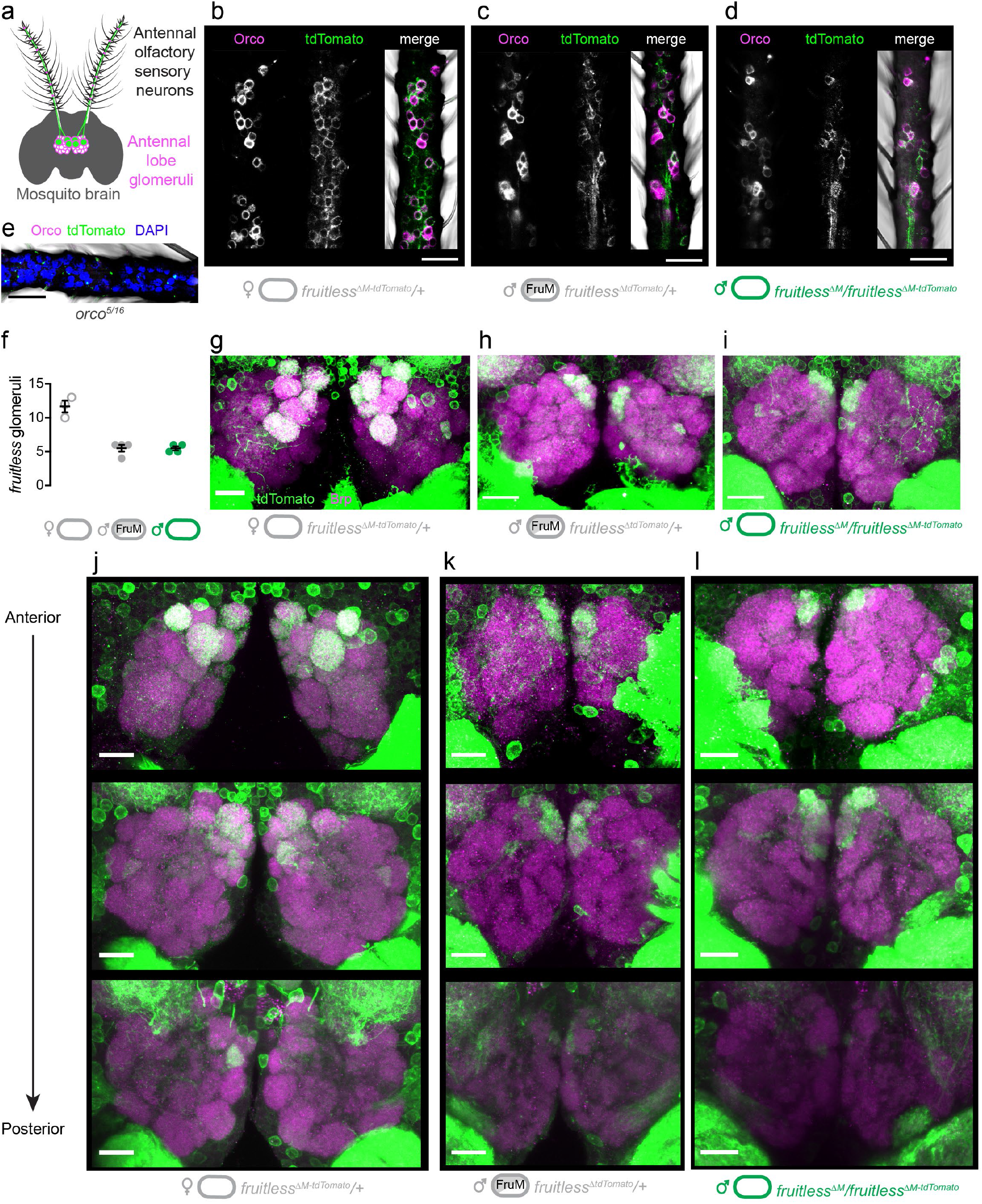
Expression of fruitless in the mosquito olfactory system. **a**, Schematic of antennal olfactory sensory neurons and their projections to the antennal lobe of the mosquito brain. **b-d**, Confocal images of antennae of the indicated genotypes with *fruitless>tdTomato* (green) and Orco (magenta) expression. **e**, Confocal image of *orco* mutant antenna, as negative control for Orco and tdTomato antibodies, with DAPI (blue). **f**, Number of antennal lobe glomeruli labeled by *fruitless>tdTomato* in the indicated genotypes. **g-i**, Confocal images of antennal lobes of the indicated genotypes with *fruitless>tdTomato* (green) and Brp (magenta) expression. **j-k**, Confocal images of antennal lobes of the indicated genotypes showing *fruitless>tdTomato* (green) and Brp (magenta) expression. Top-to bottom images are optical sections of the same lobes, arranged from anterior to posterior. All scale bars, 20 μm.

**Figure S3:**
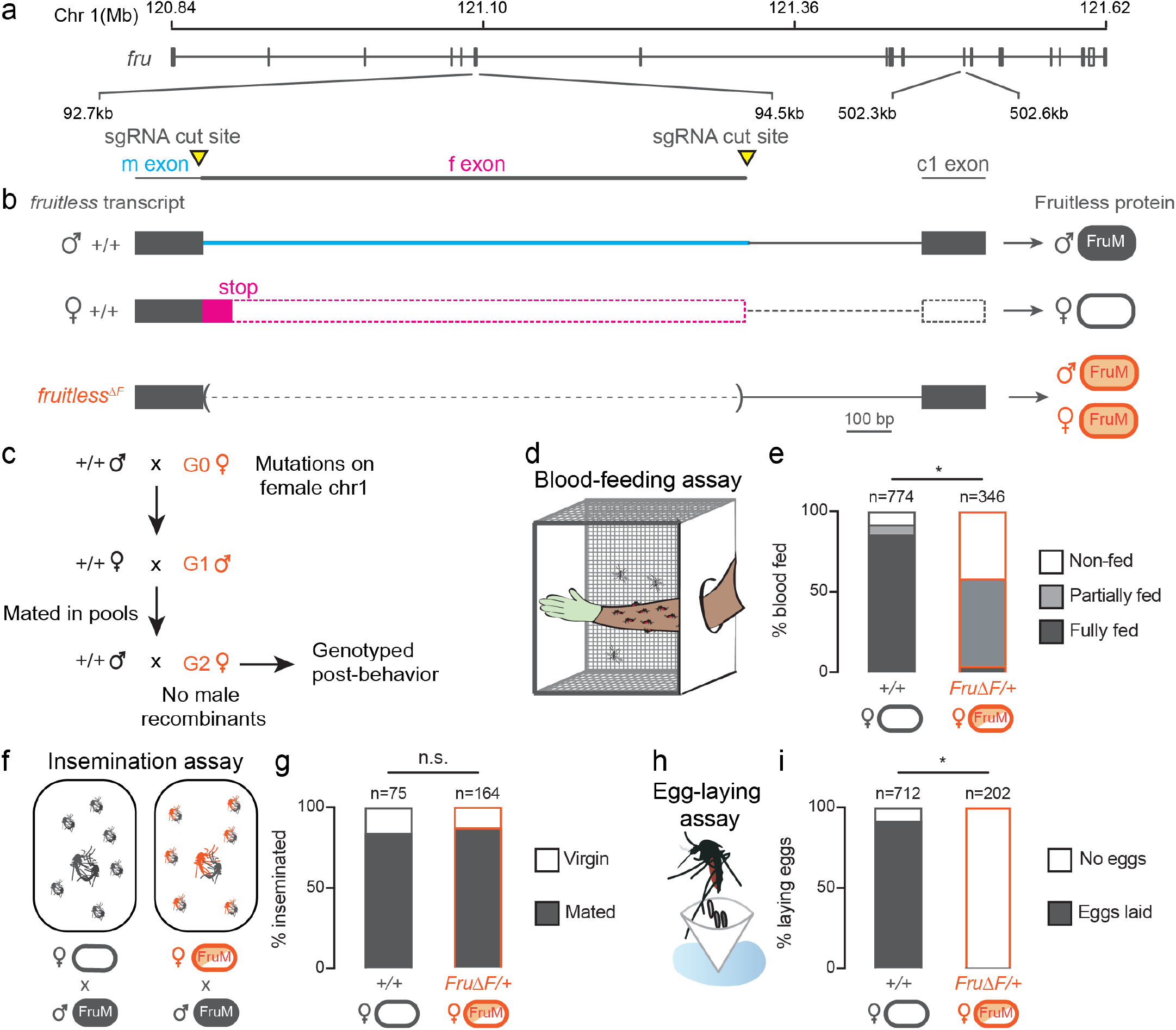
Female *fruitless*^Δ*F*^ mutant mosquitoes have blood-feeding and egg-laying defects. **a**, Schematic of *Aedes aegypti fruitless* genomic locus. **b,** Sex-specific *fruitless* transcripts and generation of *fruitless*^Δ*F*^ mutant, with effect of splicing on Fruitless protein. **b,** Crossing scheme to generate female mutants and potential male recombinants. **c**, Blood-feeding assay schematic. **e**, Feeding on live human arm; p<0.0001, Chi-square test. **f**, Insemination assay schematic. **g**, Insemination of females of indicated genotype by wild-type males; p = 0.5464, Fisher’s exact test. **h**, Egg-laying assay schematic. **i,** Egg-laying by females of indicated genotype; p<0.0001, Fisher’s exact test.

**Table S1.** List of successful and attempted genetic manipulations of the *fruitless* locus.

| Construct | Purpose | #hatched/injected<br>(% eggs hatched) | #G1<br>families<br>verified<br>hits | Hit rate<br>(%) | Notes |
| --- | --- | --- | --- | --- | --- |
| **SUCCESSFUL *FRUITLESS M* CONSTRUCTS** |  |  |  |  |  |
| *fruitlessΔM-T2A-CsChrimson-tdTomato* | Disrupt FruitlessM and express a fusion protein to label and optogenetically activate fruitless neurons in *fruitlessM* mutant males | 820/2000 (41%) | 4 | 0.97 | Viable line; tdTomato marker was bright only after antibody staining. No behavioral phenotype observed when males were exposed to red light, possibly because CsChrimson expression is low |
| *fruitlessΔM* | Disrupt FruitlessM | 376/2000 (20%) | 1 | 0.53 | Resulted from imprecise insertion of the *fruitlessΔM-T2A-QF2* construct at the fruitless locus |
| **ATTEMPTED *FRUITLESS M* CONSTRUCTS** |  |  |  |  |  |
| *fruitlessΔM-T2A-QF2* | Disrupt FruitlessM and express the bipartite Q system in fruitless neurons (regular QF2) | 376/2000 (20%) | 7 | 4.43 | All males did not mate, all females did not blood-feed |
| *fruitlessΔM-T2A-QF2w* | Disrupt FruitlessM and express the bipartite Q system in fruitless neurons (attenuated QF2) | 462/2000 (22%) | 6 | 2.61 | All males did not mate, all females did not blood-feed. In G0s crossed to QUAS-CD8-GFP, saw labeling in G1 brains that resembled *fruitlessΔM-tdTomato* expression |
| *fruitlessΔM-T2A-GCaMP6s-T2A-TdTomato-T2A-3XFLAG-DsRed-NLS* | Disrupt FruitlessM and express cytoplasmic and nuclear markers and GCaMP for calcium imaging | 349/2000 (16%) | 4 | 1.15 | Viable line but weak marker |
| **ATTEMPTED *FRUITLESS F* CONSTRUCTS** |  |  |  |  |  |
| *fruitlessΔF* | Force expression of FruitlessM in females (unmarked allele) | 205/1000 (20%) | G1s pooled for quick screening | ? | G1 males pooled and crossed to +/+ females for PCR screening. G2 females had blood-feeding defects and did not lay eggs |
| *fruitlessΔF-dsRed-3XP3* | Force expression of FruitlessM in females (marked allele) | 740/2000 (37%) | 6 | 1.60 | G1/G2 Females blood-fed normally. We suspect that the 3xP3 insertion interferes with splicing |

